# Perturbation of *in vivo* neural activity following α-Synuclein seeding in the olfactory bulb

**DOI:** 10.1101/2020.04.17.045013

**Authors:** Aishwarya S. Kulkarni, Maria del Mar Cortijo, Elizabeth R. Roberts, Tamara L. Suggs, Heather B. Stover, José I. Pena-Bravo, Jennifer A. Steiner, Kelvin C. Luk, Patrik Brundin, Daniel W. Wesson

## Abstract

**BACKGROUND:** Parkinson’s disease (PD) neuropathology is characterized by intraneuronal protein aggregates composed of misfolded α-Synuclein (α-Syn), as well as degeneration of substantia nigra dopamine neurons. Deficits in olfactory perception and aggregation of α-Syn in the olfactory bulb (OB) are observed during early stages of PD, and have been associated with the PD prodrome, before onset of the classic motor deficits. α-Syn fibrils injected into the OB of mice cause progressive propagation of α-Syn pathology throughout the olfactory system and are coupled to olfactory perceptual deficits.

**OBJECTIVE:** We hypothesized that accumulation of pathogenic α-Syn in the OB impairs neural activity in the olfactory system.

**METHODS:** To address this, we monitored spontaneous and odor-evoked local field potential dynamics in awake wild type mice simultaneously in the OB and piriform cortex (PCX) one, two, and three months following injection of pathogenic preformed α-Syn fibrils in the OB.

**RESULTS:** We detected α-Syn pathology in both the OB and PCX. We also observed that α-Syn fibril injections influenced odor-evoked activity in the OB. In particular, α-Syn fibril-injected mice displayed aberrantly high odor-evoked power in the beta spectral range. A similar change in activity was not detected in the PCX, despite high levels of α-Syn pathology.

**CONCLUSIONS:** Together, this work provides evidence that synucleinopathy impacts *in vivo* neural activity in the olfactory system at the network-level.

## Introduction

In addition to loss of substantia nigra dopamine neurons, a major pathological hallmark of Parkinson’s disease (PD) is the presence of Lewy bodies and Lewy neurites, primarily composed of insoluble misfolded aggregates of α-Synuclein (α-Syn) (Goedert, 2001; Spillantini et al., 1997). Braak and colleagues (Braak et al., 2004, 2003a), suggested that in the early stages of PD pathogenesis, α-Syn aggregates accumulate in olfactory structures, including the olfactory bulb (OB), and the enteric nervous system before appearing in other brain regions. Interestingly, ∼90% of individuals with PD exhibited olfactory deficits (Doty, 2012; Doty et al., 1988) prior to the onset of classic motor symptoms (Mahlknecht et al., 2015; Ross et al., 2008; Wu et al., 2011). Therefore, it is possible that pathogenic α-Syn aggregates within the olfactory system underlie the olfactory perceptual deficits observed in PD.

Recent studies in experimental animals have demonstrated that intracerebral inoculation of brain homogenates derived from mice and humans with synucleinopathy, or seeding recombinant pre-formed fibrils (PFFs) of α-Syn, triggers α-Syn pathology *in vivo* and *in vitro* (e.g., (Luk et al., 2012, 2009; Luk and Lee, 2014; Peelaerts et al., 2018; Rey et al., 2013). Injection of PFFs into the OB results in the spread of pathology between anatomically connected brain regions, including the piriform cortex (PCX) (Mason et al., 2016; Mezias et al., 2020; Rey et al., 2018a, 2016). Although the relationship between PD pathology and olfactory dysfunction are being explored clinically (Doty, 2017; Lee et al., 2014; Rey et al., 2018b; Wattendorf et al., 2009; Wen et al., 2017), the mechanisms underlying olfactory deficits in PD are unclear and animal modelling might provide insight into how neural processing is perturbed.

In both humans and rodents, initial odor processing occurs in the OB where olfactory receptor neurons in the nasal epithelium synapse to form OB glomeruli. Following local synaptic processing of this input (Schoppa and Urban, 2003; Wachowiak and Shipley, 2006) odor evoked information is then transferred to several secondary olfactory structures, including the PCX (Scott et al., 1980). As the primary region for processing odors, the OB is crucial for the basic initial aspects of olfaction, including the fundamental ability to detect and recognize odors. Additionally, the PCX contributes to higher-order aspects of odor perception including odor learning (Gottfried, 2010; Wilson and Sullivan, 2011). Therefore, any perturbations in odor information processing through the local neural activity of the OB and/or the PCX could result in perceptual changes (Doucette et al., 2007; Nusser et al., 2001; Wilson, 2001). Thus, accumulation of α-Syn in the OB and PCX in persons with PD (Braak et al., 2003b; Doty, 2012) and olfactory deficits observed in PD (for review (Rey et al., 2018b)) led us to propose that pathogenic α-Syn perturbs olfactory neural activity.

Prior *in situ* and *ex vivo* studies have established that the natively unfolded form of α-Syn can modulate synaptic activity (Burré, 2015; Burré et al., 2010; Chandra et al., 2004). One study used brain surface electroencephalography to uncover changes in network activity of transgenic mice overexpressing human α-Syn (Morris et al., 2015). However, no studies to date have determined whether there are effects of pathogenic α-Syn assemblies on *local* neural activity *in vivo*. Here we use the olfactory system as a model to determine the influence of α-Syn aggregates on neural activity in awake animals. Specifically, we examined spontaneous and odor-evoked local field potentials (LFPs) within the OB and PCX of mice. LFPs reflect aggregate network activity (Buzsáki et al., 2012; Mitzdorf, 1985), and in the olfactory system, LFPs are comprised of three spectral bands, including theta (2-12 Hz), beta (15-35 Hz) and gamma (40-80 Hz). These bands are thought to play unique roles in the olfaction (Kay et al., 2009). Adding to their significance, beta and gamma oscillations in key basal ganglia structures are perturbed in PD (Burciu and Vaillancourt, 2018; Little and Brown, 2014; McCarthy et al., 2011).

Here we tested the hypothesis that accumulation of pathogenic α-Syn in the olfactory system leads to aberrant LFP activity. Through multi-site LFP recordings in awake mice following α-Syn PFF seeding in the OB, we found that α-Syn PFF seeding impairs olfactory oscillatory network activity. We present evidence that synucleinopathy impacts *in vivo* neural activity in the olfactory system in manners which might contribute to the olfactory deficits associated with early PD and the prodrome of the disease.

## Materials and Methods

### Experimental subjects

A schematic of the experimental timeline is displayed in **Figure 1A**. A total of 57 female C57BL/6J mice (from donor stock originating at the Jackson Laboratory, Bar Harbor, ME) were bred and maintained in a University of Florida, Gainesville, FL vivarium. Mice were group housed until intra-cranial electrode implants as described below, with food and water available *ad libitum*. We selected female mice in order to follow the methods of previous work which characterized pathological Pser129 expression throughout the brain following OB injections of α-Syn PFFs in females (Rey et al., 2018a, 2016). While estrous stage was not monitored in the present mice, it is likely that they were in various stages of estrous on the days of recordings. Therefore, the influence of estrous cycle, if any, on the physiological measures likely averaged out. All animal procedures were approved by the University of Florida Institutional Animal Care and Use Committee and were conducted in accordance with the guidelines of the National Research Council.

**Fig 1.**
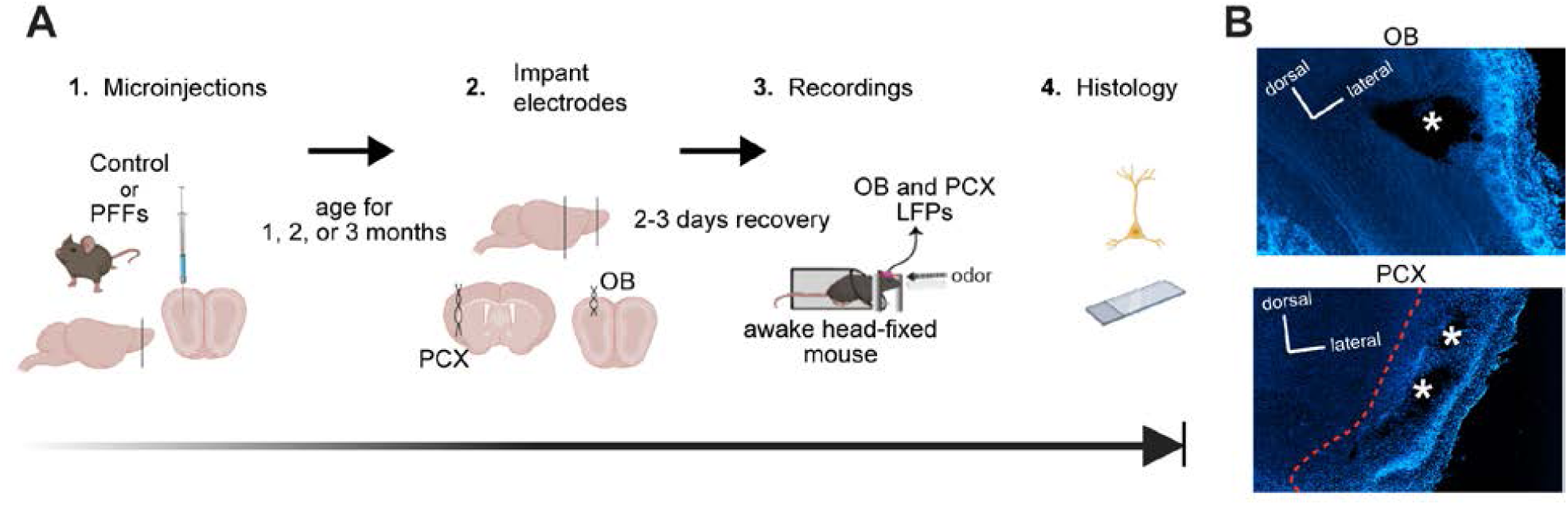
Experimental design for α-Syn seeding and subsequent multi-site LFP recordings in awake mice. **A**, 2-3 months old C57BL/6J mice received unilateral OB injections of either α-Syn PFFs or PBS and survived for 1, 2, or 3 months post injection. The mice were then surgically implanted ipsilaterally to the initial surgical site in the OB and PCX with twisted bipolar electrodes for LFP recordings. Following this, the mice were allowed 2-3 days to recover. Spontaneous and odor-evoked LFPs were recorded from awake mice while they were head fixed (to allow control for the positioning of the snout relative to the odor port) in the absence and presence of odors respectively, and then were perfused within 3 days for *post-mortem* histology. Image made in BioRender. **B**, Example localization of the electrode implants in the OB and PCX of a mouse injected with PBS using 10X magnification. * = former sites of bipolar electrode tips.

### Sonication and PFF handling

Purified recombinant full-length mouse α-Syn (as described in Volpicelli-Daley et al., 2014) was thawed at room temperature, and sonicated in a cup horn sonicator (QSonica, Q700, Newton, CT) to yield short length PFFs (**Fig. 2**). During sonication (amplitude of 50, process time of 3 mins with 1 s ON and 1 s OFF cycles), care was made to ensure the sample was submerged in water, to ensure consistent sonication power of the sample. Sonicated PFFs were stored at room temperature until being microinjected into the brain as described below.

**Fig 2.**
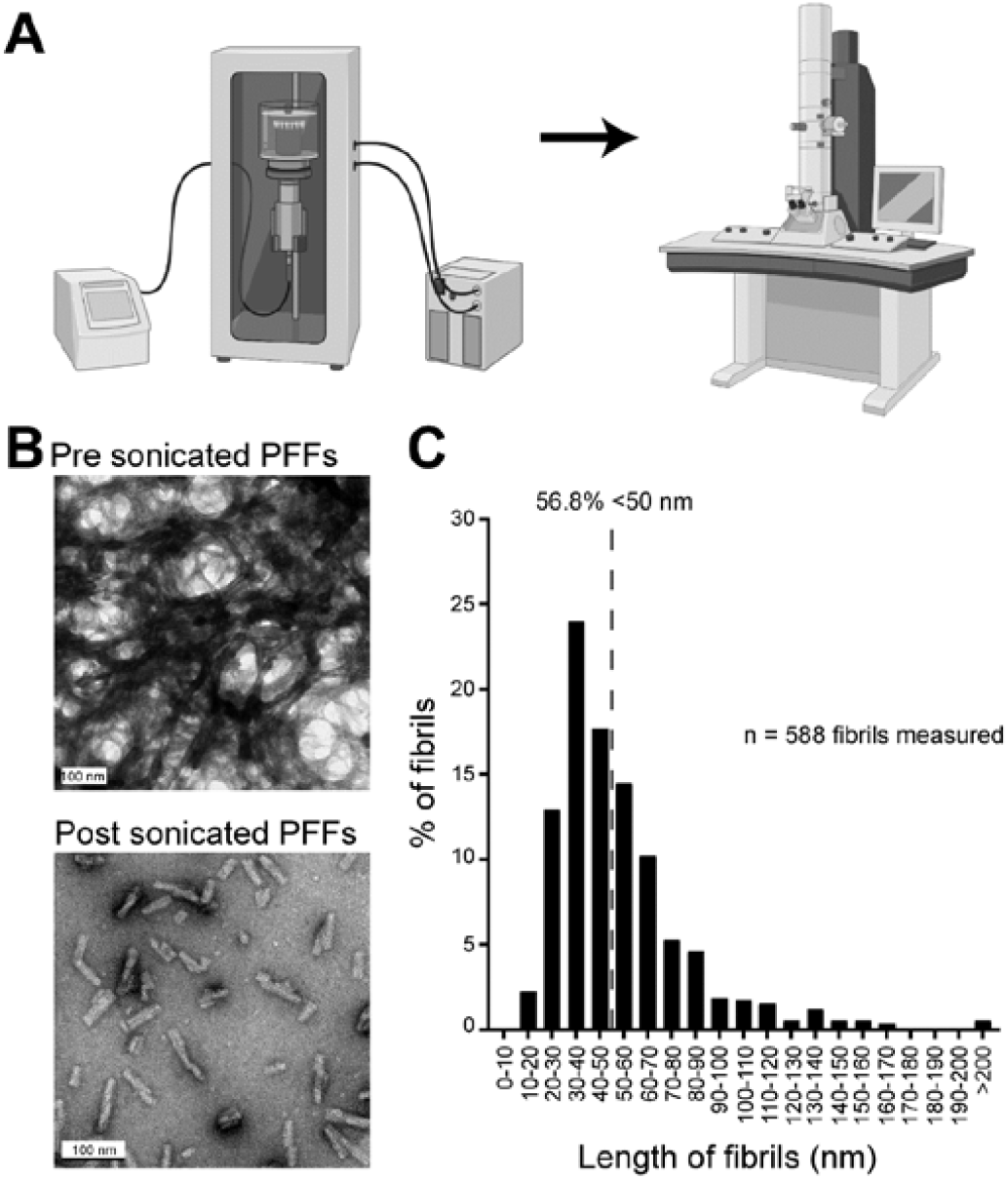
Verification of PFF length prior to OB injections. **A**, Experimental design, including sonication and electron microscopy, to optimize the length of the PFFs. Image made in BioRender. **B**, Transmission electron microscopy images of PFFs before and after sonication. To allow for visualization and quantification of sonicated individual fibrils, the sonicated sample was diluted prior to imaging. Scale bars represent 100 nm. **C**, Histogram of PFF length post sonication illustrating that >50% of the fibril population are <50 nm. Dashed line represents 50% of total population of PFFs quantified. Data from two separate sonication runs, 6-7 electron micrographs each.

### Electron microscopy

Electron microscopy was used to verify optimal PFF sonication in two separate runs of PFF samples (**Fig. 2**) following the guidelines recommended by (Polinski et al., 2018). The PFF samples from both runs were sonicated as described above. To allow for visualization and quantification of sonicated individual fibrils, the sonicated sample (not that injected into the experimental animals) was diluted (1:4) in PBS prior to imaging. The sonicated PFFs were then absorbed onto 400 mesh carbon coated grids (Electron Microscopy Sciences, Hatfield, PA) and stained with 1% uranyl acetate for subsequent electron microscopic imaging to confirm optimal sonication and thus fibril lengths. All images were captured with a Tecnai G2 Spirit TWIN 120 kV transmission electron microscope (FEI Company, Hillsboro, OR) equipped with an UltraScan 1000 (2k x 2k resolution) CCD camera (Gatan Inc., Pleasanton, CA). Fibril lengths were measured in Fiji Image J (Schindelin et al., 2012).

### Surgical procedures and animal care

#### OB microinjections

We injected PFFs unilaterally in the OB of the mice no later than 3 hours after sonication as described above. Briefly, mice were anaesthetized with isoflurane (∼3% in 1 l/min of O2) and mounted in a stereotaxic frame accompanied with a water-filled heating pad in order to maintain body temperature (38°C). Marcaine (1.7 mg/ml, s.c.; Hospira Inc., Lake Forest, IL) was injected into the site of the future wound margin. The analgesic meloxicam was also provided s.c. (5 mg/ml; Putney Inc, Overland Park, KS), and following, a midline incision was made to expose the skull cap. Next, a craniotomy (∼0.5 mm) was drilled in order to access the right OB (5.4 mm anterior to bregma, 0.75 mm lateral) and injected with either 800 nl of sterile PBS (pH 7.4; Gibco, Fisher Scientific, Hampton, NH) or 800 nl of sonicated PFFs using a glass micropipette at 2 nl/sec (1 mm ventral in the OB). Following the injection, and a resting period of 3 mins, the micropipette was gently withdrawn from the injection site at a rate of 200 μm/min. Following injections, the mice from all cohorts received *ad libitum* access to food and water, and were allowed to recover on a heating pad for at least 8 hrs.

### Implantation of LFP electrodes

Following injection of PFFs, and either 1, 2, or 3 months follow up periods, all mice were again sedated and prepared for cranial surgery as outlined above; this surgery included implantation of two pairs of twisted teflon-coated stainless steel bipolar electrodes (catalog # 791500, 0.005” diameter; A-M systems, Carlsborg, WA). The electrodes were implanted ipsilaterally to the PFF injection into both the OB and PCX. The OB coordinates were 3.8 mm anterior from bregma, 1 mm lateral, 1 mm ventral and the PCX coordinates were 0 mm bregma, 3.4 mm lateral, and 4 mm ventral. A third craniotomy was drilled over the contralateral cortex for the placement of a single stainless steel electrode to serve as the ground (catalog # 792900, 0.008” diameter, A-M Systems). This assembly was then cemented onto the skull along with a small plastic head-bar for subsequent head-fixation as defined below. After surgery, the mice were singly housed and allowed to recover on a heating pad for at least 8 hrs. These implanted mice had *ad libitum access* to food, water, and received subcutaneous meloxicam daily for 3 days (5 mg/kg).

### Outline of experimental groups

Mice were divided into two treatment groups (*i*.*e*., PBS or PFF treated) and survived for 3 different durations post injection (1, 2, or 3 months). All mice were injected at 2-3 months of age (mean age upon injection: 64 ± 1.5 days (mean ± SEM)). Total animals per group include PBS: 1 (*n*= 9), 2 (*n*=10), or 3 (*n*=8) months post injection and PFF: 1 (*n*= 8), 2 (*n*=10), or 3 (*n*=12) months post injection.

### Awake LFP recordings and data acquisition

We recorded spontaneous and odor-evoked LFP activity from 17 mice at 1 month post injection (*n*= 9 PBS, *n*= 8 PFF), 20 mice at 2 months post injection (*n*= 10 PBS, *n*= 10 PFF), and 20 mice at 3 months post injection (*n*= 8 PBS, *n*= 12 PFF). During any given day, mice of more than one condition [1, 2, or 3 months post injection and/or PBS/PFF treatment group] were used in recordings. A schematic of the recording session structure is shown in **Figures 3A & 3B**. All the recordings were performed in a dimly-lit, well-ventilated room maintained at 20-22°C, between 0900 and 1800 hrs.

**Fig 3.**
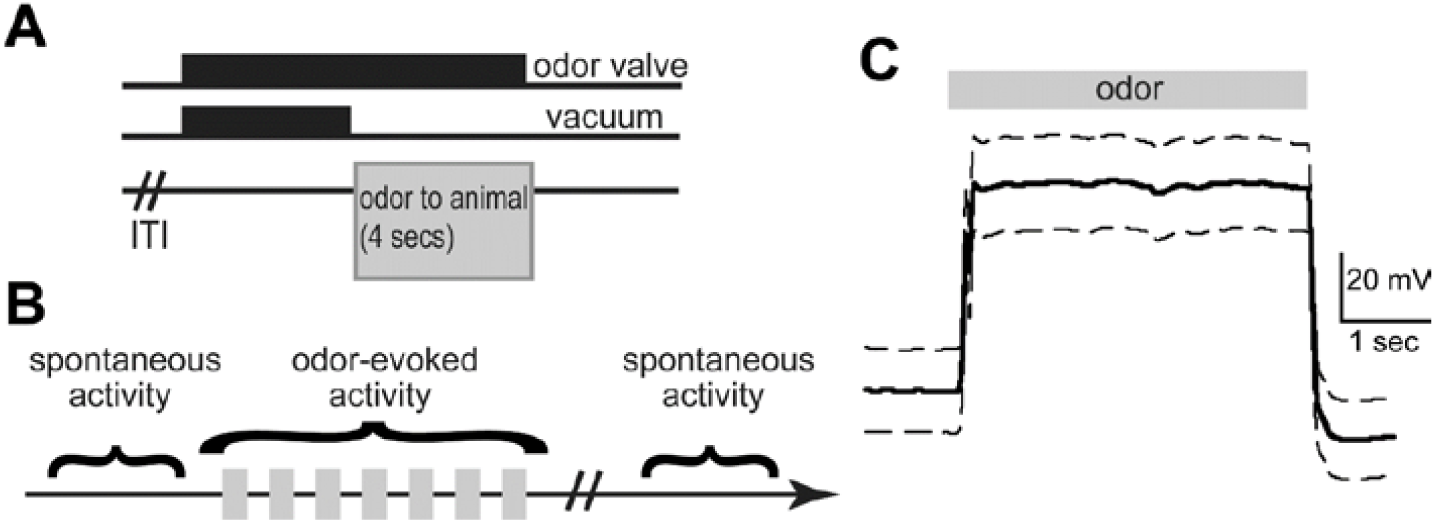
Paradigm for recording spontaneous and odor-evoked LFPs from head-fixed awake mice. **A**, After a variable inter-trial interval (ITI), the odor valves are turned on for 8 secs and the vacuum for 4 secs. This allows the animal to be presented with an odor for 4 secs. **B**, Schematic showing the recording paradigm. An epoch of spontaneous LFP activity was recorded before and after odor presentation. During odor presentation, 7 odors were presented in a pseudo-random order for 4-5 sessions, and their odor-evoked activity recorded. **C**, Average photoionization detector trace in response to 12 presentations of 1 Torr isopentyl acetate, depicting the rapid temporal dynamics and stability of odor presentation (10 Hz, low pass filtered). Data are mean +/-SEM.

Mice were head-fixed and an odor-port was positioned ∼2 cm from the nose, prior to the start of the recordings, as we have described previously (Gadziola et al., 2015). To monitor spontaneous LFP activity, we allowed the animal to rest during head-fixation for several minutes (∼5 mins) prior to and following a series of odor deliveries (**Fig. 3B**). While optimally, animals would have been habituated to this paradigm for days to mitigate stress which may influence neural activity, in order to reduce variability within groups, we sought to strictly schedule and record from each animal on a single day at a precise time post injection. All odors and the blank stimulus (mineral oil) were presented for 4 secs each, in a semi-automated pseudorandom order for a total of 4 times each, with an approximately 15 secs inter-stimulus interval. Throughout recordings, OB and PCX activity was acquired at 2 kHz and filtered (100 Hz, 2^nd^ order low-pass, 60 Hz notch) along with stimulus presentation events using a Tucker Davis Technologies RZ5D amplifier (Alachua, FL).

### Stimulus delivery

For odor presentation, odors including isopentyl acetate, 2-butanone, and 1,7-octadiene (Sigma Aldrich, St. Louis, MO) were each diluted in their liquid state to 133.332 Pa (1 Torr) and 266.645 Pa (2 Torr) in light mineral oil (Sigma Aldrich) which also served as the blank stimulus. Stimulus vapors controlled with an air-dilution olfactometer were run from glass headspace vials (100 ml/min) where they were later blended with clean nitrogen (900 ml/min) in the odor port thereby yielding a total odor flow rate of 1 L/min. The olfactometer was equipped with independent stimulus lines up to the point of entry into a Teflon odor port, in order to eliminate chances of cross-contamination of the stimuli and also to allow for rapid temporal control of odor dynamics as they reach the animal. To confirm the dynamics of the odor plume as it leaves the odor port, we used a photoionization detector (Aurora Scientific, Aurora CO). As shown in **Figure 3C**, odor delivery occurred rapidly, and was largely stable throughout the 4 sec of delivery.

### Tissue collection and histology

Within 48 hrs after recordings, the mice were overdosed with Fatal-plus (0.01 ml/g; Vortech Pharmaceutical Ltd, Dearborn, MI). Following confirmation of deep sedation, they were perfused with cold saline and subsequently 10% phosphate buffered formalin. Brains were collected and stored at 4 °C in 10% formalin / 30% sucrose prior to sectioning. Serial 40 µm thick coronal sections were collected using a sliding microtome, and stored in Tris-buffered saline (TBS, pH 7.4) with 0.03% sodium azide. These sections were used for electrode verification and phosphoserine 129 (Pser129) immunofluorescence.

### Pser129 immunofluorescence

Presence of Pser129 α-Syn-immunopositive inclusions is considered pathological given the abundance of this post-translationally modified form of α-Syn in Lewy pathology (Fujiwara et al., 2002). Our previous work has also shown that Pser129 α-Syn-immunopositive inclusions in the animal model we used in the present study are positive for ubiquitin, p62, Thioflavin S and are Proteinase K resistant, all features of clinical Lewy pathology (Rey et al., 2016). Thus, we used Pser129 immunofluorescence to quantify pathological burden resultant from OB PFF or PBS microinjections (the latter as a control). Floating OB and PCX sections from PFF and PBS injected mice were rinsed thrice in TBS and subsequently, dilution buffer (10 mins each). The dilution buffer was comprised of 2% bovine serum albumin (Sigma Aldrich), 0.9% sodium chloride (Sigma Aldrich), 0.4% Triton-X 100 (Sigma Aldrich), and 1% normal goat serum (Sigma Aldrich) in TBS. Next, the sections were blocked in 20% normal donkey serum (Sigma Aldrich) in TBS for 20 mins and incubated for 24 hrs in primary antibody rabbit anti-Pser129 (1:5000 in dilution buffer, catalog # EP1536Y, Abcam, Cambridge, MA) at room temperature. On the following day, all sections were rinsed four times in dilution buffer (10 mins) and incubated for 2 hrs in secondary antibody Alexa Fluor 594 goat anti-rabbit IgG (1:500 in dilution buffer, catalog # A11012, Invitrogen, Carlsbad, CA). Sections were rinsed thrice in TBS and twice in double distilled water (5 mins each). Finally, tissue was placed on slides and cover-slipped with mounting media containing DAPI (Fluoromount-G; 4’,6-diamidino-2-phenylindole; Invitrogen). All immunofluorescence runs contained tissue from more than one age (1, 2, or 3 months post injection) and treatment group (PBS or PFF).

### Electrode verification

PCX electrode tips/recording sites were verified with microscopy. Due to the ease of targeting, not all mice used in the study underwent exclusive examination for electrode tip placement in the OB. For PCX recording site verification, tissue processed for anti-Pser129 immunofluorescence (counterstained with DAPI) was used. When additional sections were needed for confirmation of recording sites, additional PCX tissue was placed on slides and counter-stained with DAPI. Only one mouse (PBS, 2 months post injection) did not contribute PCX data since the electrode was misplaced. The total numbers of animals contributing data / brain region is defined below. Representative images of electrode tips localized in the OB and separately in the PCX are displayed in **Figure 1B**.

### Pser129 imaging and quantification

The OB and PCX were identified based on an established brain atlas (Paxinos and Franklin, 2000). Images of the OB and PCX were acquired from the hemisphere ipsilateral (identified from the electrode tracks) to the OB microinjection. All Pser129 imaging was performed with a Nikon Ti2e microscope equipped with a 15MP monochrome camera and using 20X magnification. Additionally, image acquisition settings, particularly the gain, light intensity, and exposure were kept constant for all images. First, images were collected which included the OB and PCX (≥4 images/region). Attempts were made to acquire images from similar anterior-posterior extents of each brain area. Next, by a treatment group blind experimenter, the images were cropped so that the resultant image included solely OB and PCX, and that the images spanned the majority of cell layers, as exemplified in **Figure 4**. Also by a treatment group blind experimenter, these cropped images were next thresholded using semi-automated routines in Nikon Elements and with fixed settings across all images for later quantification of Pser129 levels. The % area in each of the cropped and thresholded images occupied by Pser129 was then calculated as a ratio of the pixel area above threshold : the total pixel area. Representative images showing Pser129 burden in the OB and PCX are presented in an inverted grayscale (**Fig. 4B**), to allow ease of visualization of puncta, although all of the data analysis steps outlined above occurred in the original high-resolution color images.

**Fig 4.**
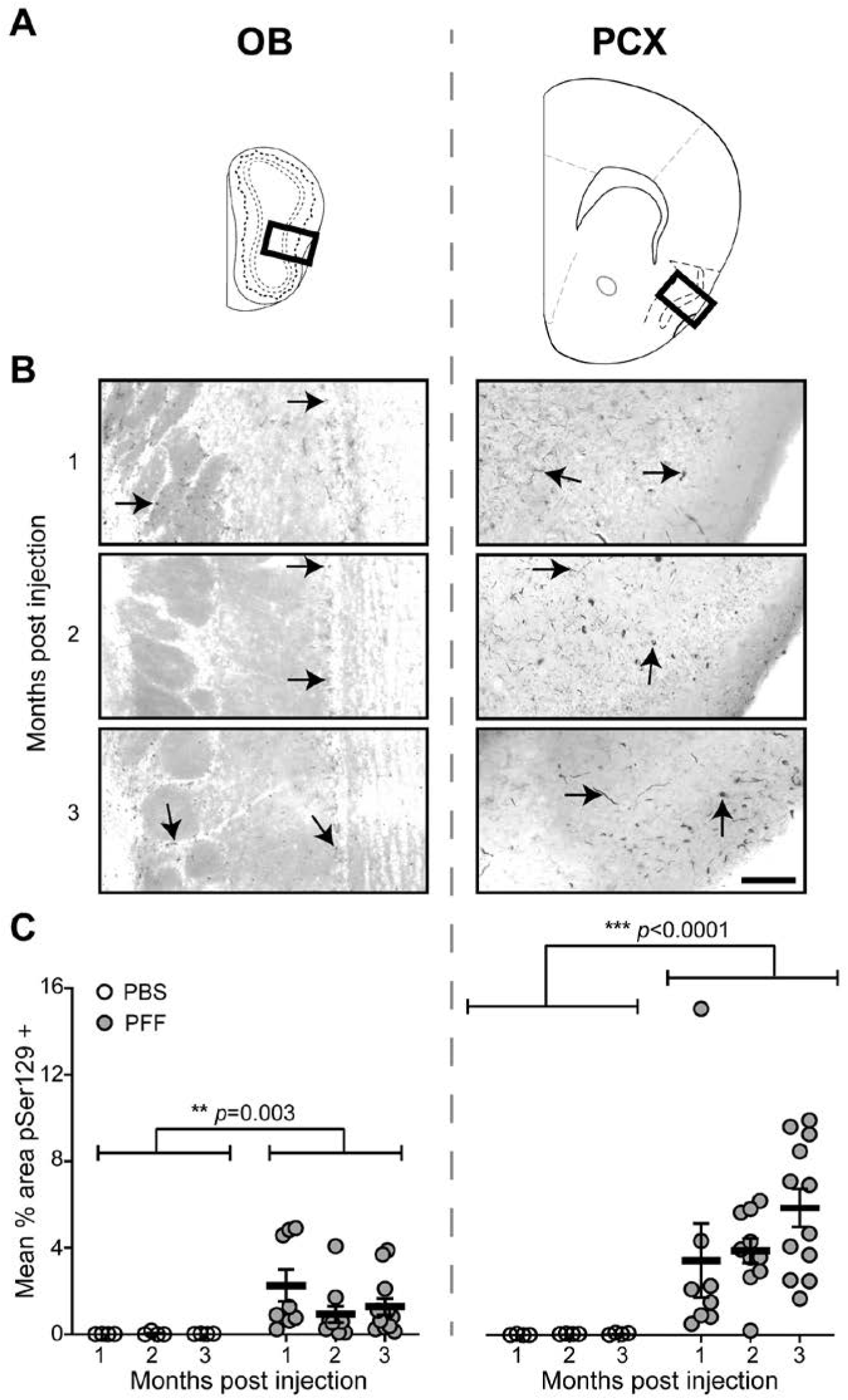
PFFs injected in the OB induced an amplification and spread of pathology to interconnected regions, including the PCX. Pser129 immunofluorescence was used as an assay to detect pathological α-Syn. **A**, Coronal panel showing the regions (bold boxes) used for quantifying Pser129 expression level in the OB and PCX. **B**, Representative images of Pser129 immunofluorescence staining of the OB and PCX of mice that survived for 1, 2, or 3 months post PFF seeding. Arrowheads in the OB and PCX panels indicate areas of neuritic pathology. The images were gray-scaled and inverted to show pathology more readily, for illustration purposes of this figure only. **C**, Quantification of mean % area in the OB and PCX, showing Pser129 immunofluorescence in PFF and a subset of PBS injected mice that survived for 1 (PFF *n*= 8, PBS *n*= 4), 2 (PFF *n*= 10, PBS *n*= 4), and 3 (PFF *n*= 12, PBS *n*= 4) months post injection. Animals injected with PFFs had a significantly greater Pser129 immunopositive signal than the PBS injected animals, including in both the OB and PCX. Significant increase in mean % area Pser129 was observed in the PCX when compared to the OB. \*\*\**p* ≤ 0.001 ANOVA followed by Tukey’s multiple comparison’s test.

### Analysis and Statistics

In total, we acquired spontaneous and odor-evoked LFP activity from 17 mice at 1 month post injection (*n*= 9 PBS, *n*= 8 PFF), 20 mice at 2 months post injection (*n*= 9 PBS, *n*= 10 PFF), and 20 mice at 3 months post injection (*n*= 8 PBS, *n*= 12 PFF). As described in the electrode verification section above, these numbers only include mice which had electrodes verified in each of the target brain regions. All of these mice contributed clean artifact free signals and also passed the electrode placement confirmation described above.

Analysis of the LFP data was performed by experimenters blinded to the experimental groups using Spike 2 (Cambridge Electronic Design, Milton, Cambridge). Data were processed using Fast Fourier Transform (FFT) analysis [for spontaneous (1.006 secs Hanning window and 0.994 Hz resolution) and odor evoked (0.503 sec Hanning window and 1.987 Hz resolution)] in order to classify and distribute the data within different LFP frequency bands. While due to the time-window sizes, we used differing FFT resolutions for the above calculations, no comparisons are made between the spontaneous and odor-evoked events. Power spectra of spontaneous and odor-evoked LFP activity were extracted from 0-100 Hz. LFP spectral bands were defined as theta (1-10 Hz), beta (10-35 Hz), and gamma (40-70 Hz) (Kay et al., 2009). OB-PCX spontaneous LFP activity was analyzed over a duration of 200 secs sampled both prior to and following presentation of odors (at the start and end of the recording session respectively). Odor-evoked response magnitude was used as a variable to measure odor-evoked LFP activity. OB-PCX odor-evoked response magnitude is a ratio of time matched (4 secs) LFP power during odor presentation to LFP power before odor presentation. Herein we focus our odor-evoked analyses solely on 1 Torr isopentyl acetate trials. This is based upon the inconsistent responses elicited by 2-butanone and 1,7-octadiene in all mice, including those PBS treated. Further, when collapsing across groups, we did not find an effect of 1 Torr versus 2 Torr intensity of isopentyl acetate and so for simplicity we restrict all analyses to 1 Torr isopentyl acetate.

All statistical analyses were performed using Microsoft Excel and Graphpad Prism 8 (San Diego, CA). Any comparison between PFF and PBS injected mice or between brain regions was conducted using 2-way ANOVA. Mixed effects analyses were used to test for possible time-post injection effects (1, 2, or 3 months post injection). Post-hoc analyses included multiple comparison corrections whenever appropriate. All data are reported as mean ± SEM unless otherwise indicated.

## Results

### Accumulation of Pser129 pathology in key olfactory structures following OB α-Syn seeding

To induce α-synucleinopathy in the olfactory system, PFFs or PBS as a control, were unilaterally injected into the OB of 2-3 months-old female mice. The mice were then allowed either 1, 2, or 3 months time post-injection prior to being implanted with electrodes into their OB and PCX, ipsilateral to the injection, and LFP recordings performed (**Fig. 1**) as described below. Following recordings, the brains were collected and processed for anti-Pser129 immunofluorescence. Confirming previous results using this OB PFF seeding model (Rey et al., 2018a, 2016), we observed Pser129 burden in key olfactory structures, including the OB itself and the PCX, in mice injected with PFFs (**Fig. 4**). Pser129 immunostaining was detected throughout all cell layers in both brain regions of PFF injected mice (**Fig. 4B**). There was a significant effect of PFF treatment on Pser129 levels in the OB (*F*(1,36) = 10.56, *p* = 0.003), as well as the PCX (*F*(1,36) = 19.84, *p* < 0.0001) (**Fig. 4C**). Additionally, across all time groups (1 to 3 mo), there was an effect of brain region on Pser129 immunostaining indicating that the PCX was more vulnerable to accumulation of misfolded α-Syn compared to the OB (*F*(1, 22) = 16.63, *p* = 0.0005). There was a non-significant trend towards increased levels of Pser129 in the PCX in the 3 versus the 1 month group (*F*(2,27) = 1.611, *p* = 0.218) which is in contrast to the largely stable Pser129 levels within the OB from 1 to 3 months (**Fig. 4C**).

### Preservation of spontaneous LFP dynamics following α-Syn seeding

We began our investigation into the influence of α-Syn aggregates on LFP dynamics by analyzing spontaneous (*viz*., those in the absence of experimentally-applied odors) levels of theta, beta, and gamma-band powers in the OB and PCX of mice that survived 1, 2, or 3 months post injection (the same mice used for histology above). As is well established (for review (Kay et al., 2009)), spontaneous OB and PCX LFP activity is characterized by prominent theta band power coupled to the respiratory cycles, along with beta and gamma band power which also occur in somewhat phasic manners with respiratory theta (**Fig. 5A & 5B**). Since the full spectrum is diverse in power (**Fig. 5B**), as is standard when quantifying olfactory LFPs, throughout this paper we calculated the power of each of these spectral bands separately and focused on testing for PFF treatment effects by comparing PBS vs PFF treated animals.

**Fig 5.**
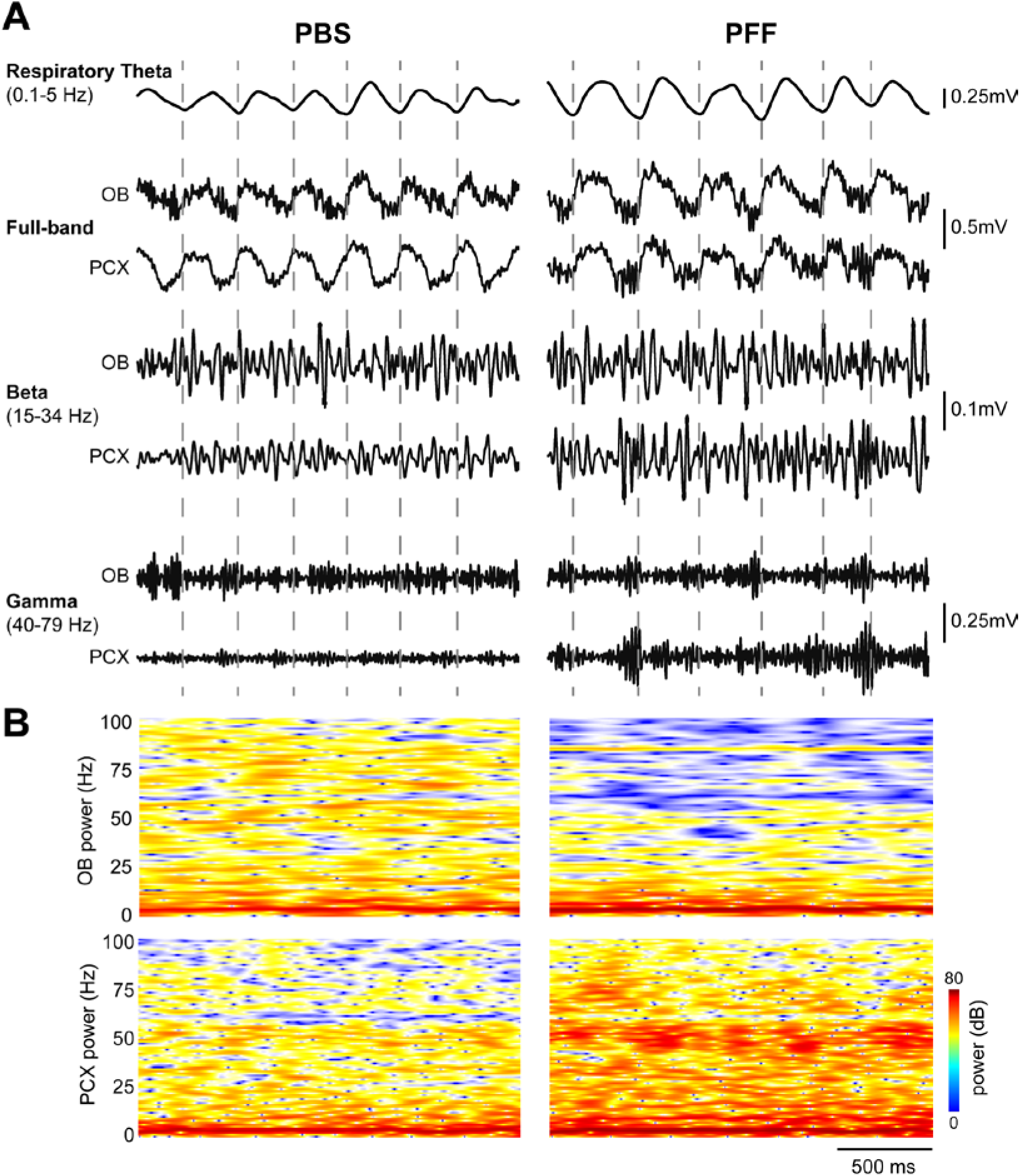
Example spontaneous LFP activity. **A**, Representative spontaneous LFP traces from two separate mice injected with either PBS (left) or PFF (right), 2 months prior to recording. Shown are full band traces from simultaneous OB and PCX recordings (0.1-100Hz) which were also filtered to separately display beta and gamma band activity as defined in the figure. Respiratory theta from the OB is also displayed along with dashed vertical lines indicating the timing of OB respiratory cycles for visual aid. **B**, 2-dimensional histograms of the same spontaneous full band power spectrograms with power displayed in dB to help illustrate the diversity of the full band data.

Despite the significant Pser129 burden in each structure (**Fig. 4**), we did not find an overall (across age groups) effect of PFF treatment on spontaneous activity in either the OB or PCX (**Fig. 6**). In the OB specifically, there was no change in theta (*F*(1,51) = 0.993, *p* = 0.324), beta (*F*(1,51) = 0.035, *p* = 0.853), or gamma band powers (*F*(1,51) = 0.013, *p* = 0.910) when comparing PBS injected animals to those injected with PFFs (**Fig. 6A**). Similarly in the PCX, we did not find an overall change in theta (*F*(1, 50) = 0.017, *p* = 0.897), beta (*F*(1, 50) = 0.084, *p* = 0.774), or gamma band powers (*F*(1, 50) = 0.073, *p* = 0.788) in PBS versus PFF injected animals (**Fig. 6B**). Thus, despite pathogenic propagation of α-Syn from the seeded structure (the OB) into a monosynaptically interconnected structure (the PCX), the spontaneous LFP activity in neither brain region was significantly disrupted.

**Fig 6.**
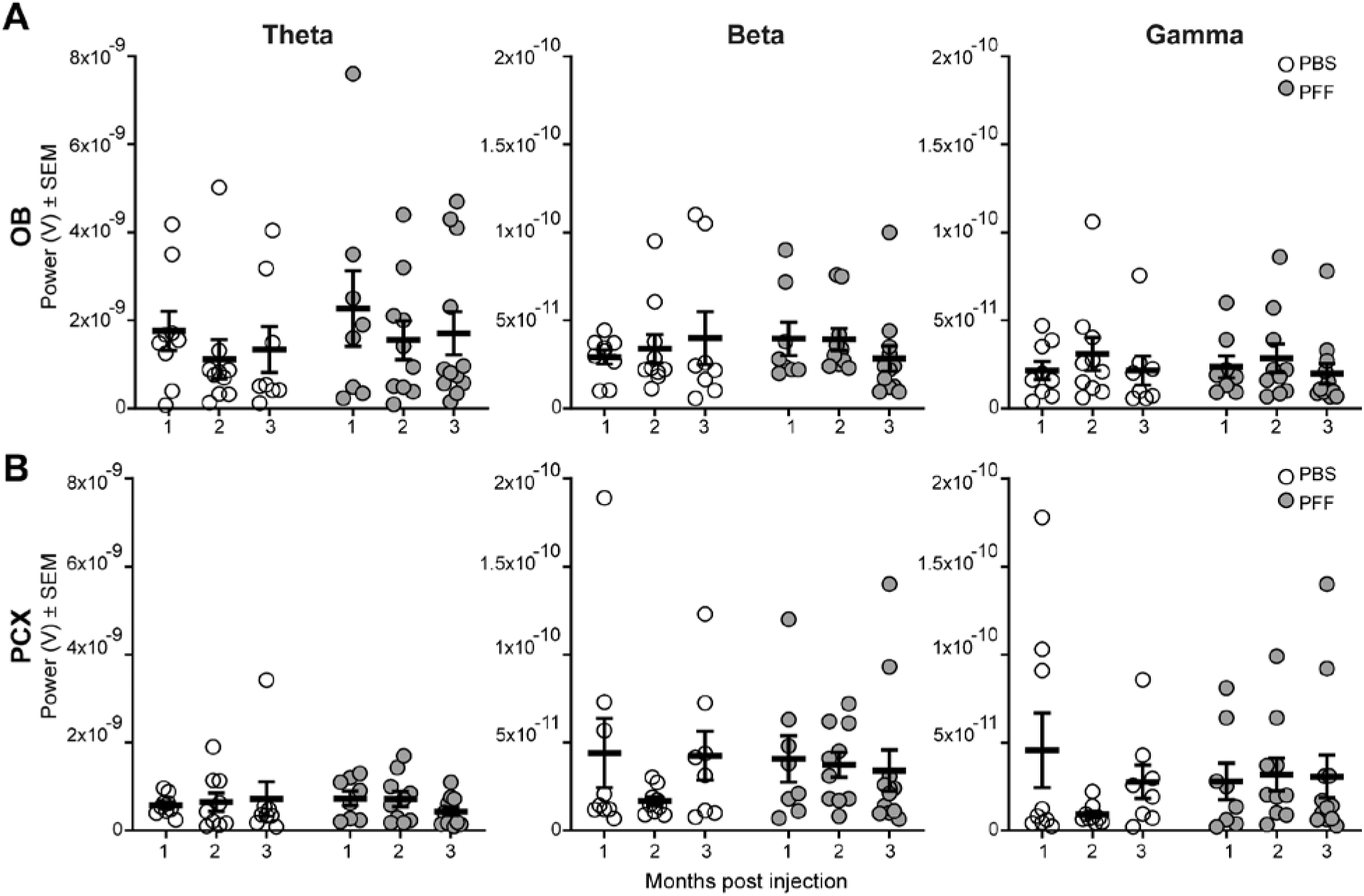
No effect of PFF injection on spontaneous LFP powers in the OB or PCX. Spontaneous OB (**A**) and PCX LFP power (**B**), consisting of theta (1-10 Hz), beta (15-34 Hz), and gamma (40-75 Hz) spectral bands in either PFF seeded mice or PBS injected mice that survived for 1 (PFF: *n*= 8, PBS: *n*= 9), 2 (PFF: *n*= 10, PBS: *n*= 9), or 3 (PFF: *n*=12, PBS: *n*=8) months post injection. No treatment or age-post injection effect was observed in either brain region.

### α-Syn seeding results in heightened OB beta-band odor-evoked activity

The OB and PCX are important for the formation of olfactory perception, and the processing of odor information in each of these stages is considered critical for olfaction. Odor-input drives increases in LFP activity in the OB (Kay et al., 2009). Beta band power is especially considered important for the coordinated transfer of sensory information between brain regions (Haegens and Zion Golumbic, 2018; Kopell et al., 2000) and thus any changes in beta activity during odor may be especially influential to perception. Therefore, we next took advantage of the monosynaptic and bisynaptic inputs of odor information into these structures by analyzing odor-evoked LFPs (**Fig. 7**) and how OB PFF seeding may impact odor-evoked activity. As expected based upon a wealth of prior research, odor-evoked increases in LFP power were observed in both the OB and PCX of PBS treated mice **(Fig. 7**, left panels). These example traces also suggested that the power may be different during odor in the PFF treated animals compared to those treated with PBS. To directly compare between groups, across all trials of the odor (1 Torr isopentyl acetate), we calculated the LFP spectral power during the 4 seconds odor was presented to the mouse’s nose and also the 4 seconds immediately prior to odor delivery. We then computed the ratio of the during odor epoch : the pre-odor epoch, in order to calculate an odor-evoked power ratio. We then averaged the calculated ratios across all trials in the session (4-5 trials/animal) to calculate each animal’s average odor-evoked power ratios as a simple measure of odor-evoked network activity (**Fig. 8**).

**Fig 7.**
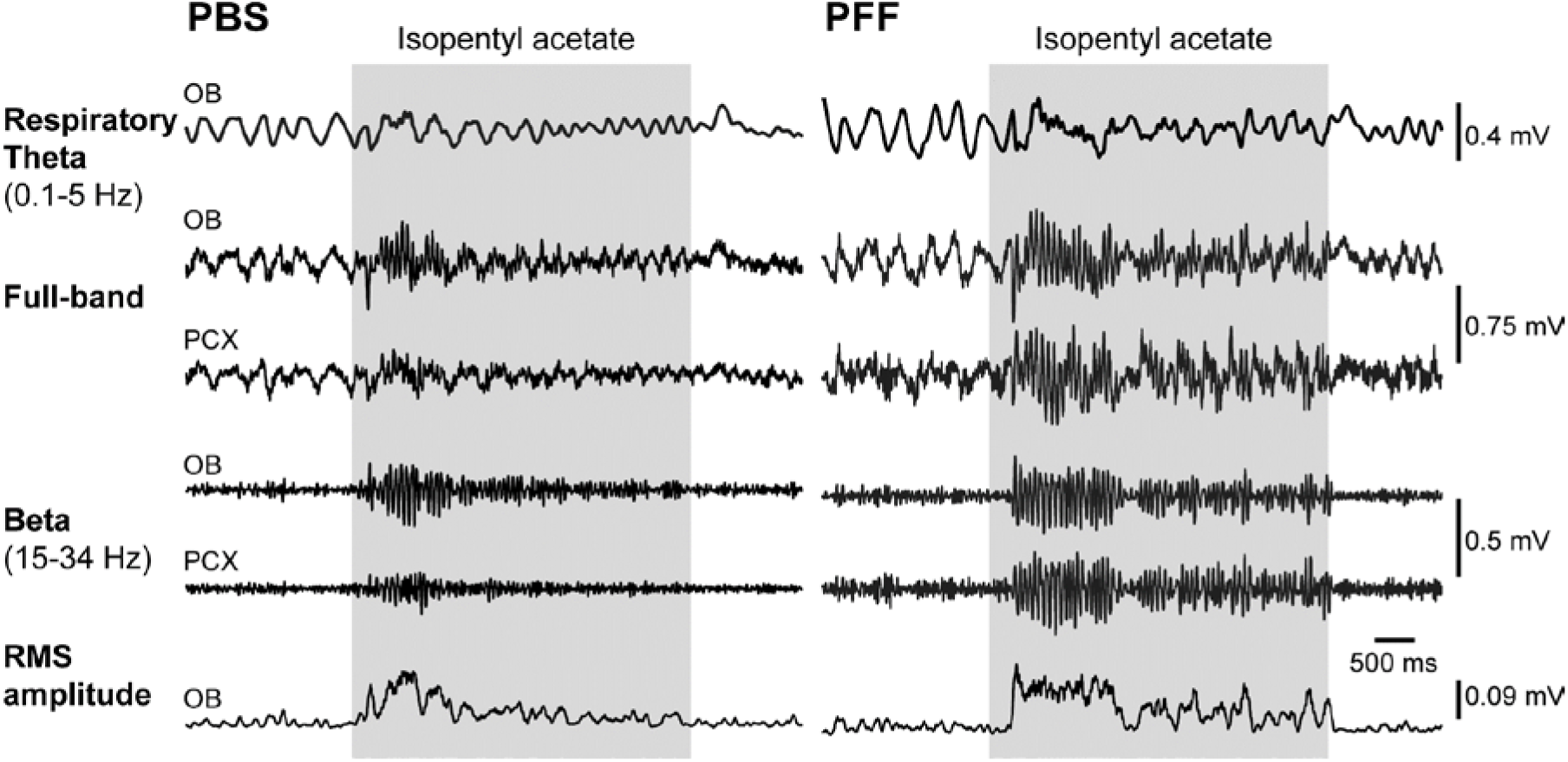
OB PFF seeding entails heightened odor-evoked OB beta-band power. Representative odor-evoked LFP traces from two separate mice injected with either PBS (left) or PFF (right), 2 months prior to recording. Shown are full band traces from odor-evoked OB and PCX recordings (0.1-100Hz) which were also filtered to separately display beta band activity as defined in the figure. Respiratory theta from the OB is also displayed as is the root mean square of the beta band activity to illustrate elevated power of beta activity in the PFF injected versus PBS injected mouse. Gray shaded boxes indicate the time of odor delivery.

**Fig 8.**
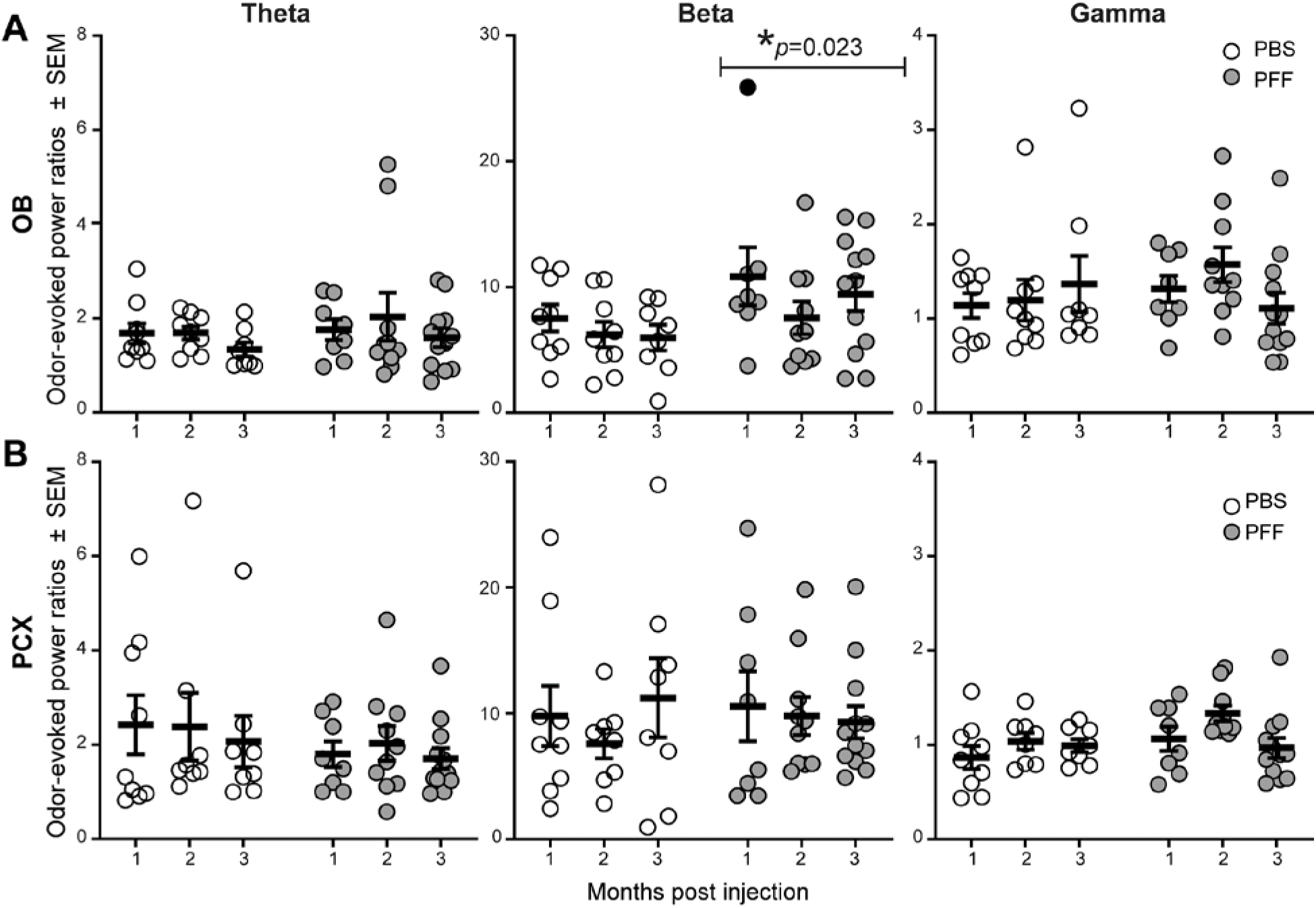
Analyses of odor-evoked activity uncover PFF seeding induced aberrant OB beta-band power during odor. **A**,**B**, Odor-evoked OB and PCX LFP power, consisting of theta (1-10 Hz), beta (15-34 Hz), and gamma (40-75 Hz) spectral bands in either PFF seeded mice or PBS injected mice that survived 1 (PFF: *n*= 8, PBS: *n*= 9), 2 (PFF: *n*= 10, PBS: *n*= 9), or 3 (PFF: *n*=12, PBS: *n*=8) months post injection. Across all age groups, a significant increase in the beta band power in the OB of PFF seeded mice was observed when compared to PBS treated. *ANOVA. Solid data point indicates an animal whose elevated OB beta power was associated with high OB Pser129 pathological burden.

In the OB, while analyses of odor-evoked power ratios did not uncover an effect in either the theta (*F*(1, 50) = 0.944, *p* = 0.336) or gamma bands (*F*(1, 50) = 0.389, *p* = 0.536), there was a significant effect of PFF treatment on increasing odor-evoked beta band power (*F*(1, 50) = 5.531, *p* = 0.023) (**Fig. 8A**). Post-hoc tests within these spectral bands, and also within individual age groups, did not reveal a similar effect of PFF treatment (*p >* 0.05, Sidak’s test for correction of multiple comparisons). Nevertheless, there is a clear elevation in beta-band odor evoked power ratios in the PFF-treated animals, which is comparable across the groups from 1 to 3 months post PFF injection.

In contrast to the OB, theta (*F*(1, 49) = 1.351, *p* = 0.251), beta (*F*(1, 49) = 0.042, *p* = 0.839), and gamma band odor-evoked powers (*F*(1, 49) = 3.380, *p* = 0.072) were similar between PBS and PFF treated animals across all ages in the PCX (**Fig. 8B**). This indicates that, at least at the level of LFP monitoring, the effect of α-Syn OB seeding is most dramatic in shaping OB odor-evoked activity. This is a significant finding since the OB provides nasal-derived odor information into the entirety of down-stream brain regions.

### Are the changes in beta-band dynamics correlated with Pser129 levels?

Finally, we analyzed whether the levels of Pser129 immunostaining within the OB were correlated with the above levels in odor-evoked beta-band power. We focused on this possible relationship since the odor-evoked beta-band activity was significantly elevated in the OB following PFF injection (**Fig. 8**). Further, some PFF-treated animals contributed abnormally high levels of odor-evoked beta-band activity (**Fig. 8**, dark data point). Indeed, upon inspection, we determined that some of these mice were also those which displayed elevated Pser129 pathological burden in the olfactory system and that led us to predict these factors were related. Surprisingly, no significant correlation was found between levels of OB Pser129 and odor-evoked power ratios within beta activity in the OB (Pearson *r*(28) = 0.156, *p* = 0.411) (**Fig. 9**). Since, as discussed earlier, beta band activity in the OB may originate from centrifugal input coming from the PCX, we also tested whether a correlation exists between OB odor-evoked beta power and PCX Pser129 levels, yet also found no significance (Pearson *r*(28) = -0.018, *p* = 0.923) (not shown). Thus, the aberrant odor-evoked beta band activity observed in PFF treated mice is not correlated with levels of Pser129 pathology.

**Fig 9.**
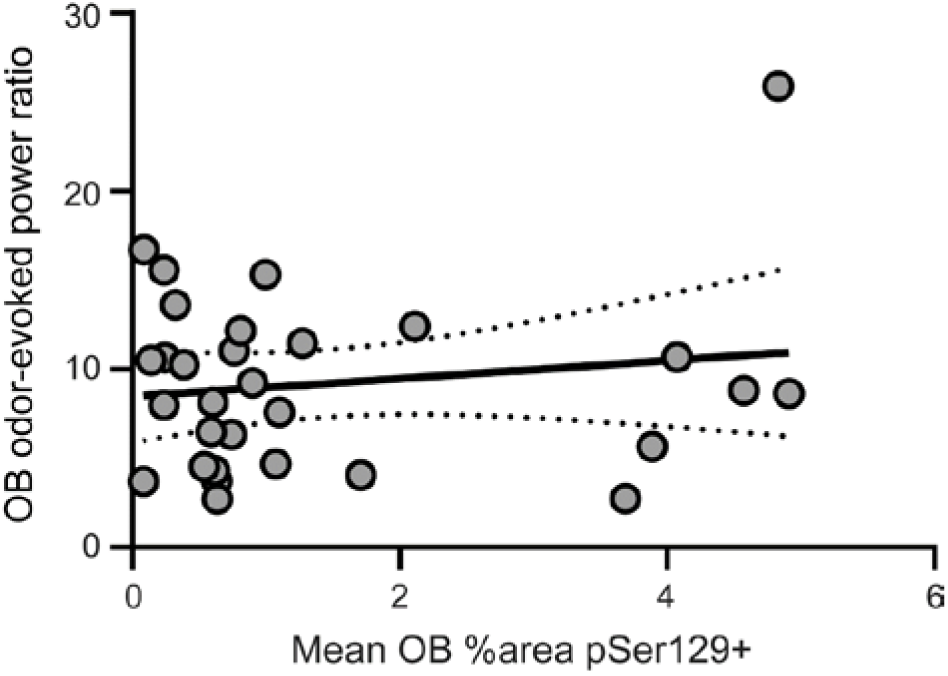
Lack of statistical correlation between OB Pser129 burden and aberrant beta band activity. Scatterplot illustrating the relationship between the mean % area occupied by Pser129 in the OB (as quantified in **Fig. 4**) and odor-evoked beta power (gray, 15-34 Hz). Gray dashed line indicates the linear fit bounded by 95% confidence intervals (Pearson *r*(28) = 0.156, *p* = 0.411).

## Discussion

We tested the hypothesis that progressive development of α-Syn aggregates in the olfactory system impacts neural dynamics. While key regions within the basal ganglia system display aberrant synchrony of neural dynamics which may contribute to motor deficits in PD (Brown and Williams, 2005; De Hemptinne et al., 2015; Hammond et al., 2007; Kühn et al., 2006; Moran et al., 2008), the direct contributions of α-Syn aggregates to neural dynamics *in vivo* are unresolved. This is an important question because hyposmia is prevalent in PD, and often develops before the onset of clear motor deficits. Furthermore, greater insight into the impact of α-Syn aggregates on neural circuits is important for our understanding of functional deficits in PD and related synucleinopathies. Perturbations in neural synchrony might be highly relevant to the symptomology of clinical synucleinopathies since they may alter key aspects of function, from cognitive, motor, to sensory – depending upon the system impaired.

We used α-Syn seeding in mice in combination with olfactory LFP recordings. The olfactory system is an excellent model to test whether pathogenic α-Syn impacts neural dynamics, given both the known aggregation of Lewy bodies in the OB during the earliest states of PD, and decades of work carefully examining the dynamics of LFPs in the OB and PCX, including how these dynamics are influenced by odors. Importantly, LFPs are reflective of aggregated network activity (Buzsáki et al., 2012; Mitzdorf, 1985) and are considered a substrate for the rhythmic sampling of sensory information (Haegens and Zion Golumbic, 2018). Therefore, changes in LFPs may impact odor perception. Here we found that the progressive development of α-Syn aggregates influenced specific sensory-evoked LFP activity in region-selective manners (*viz*., not all brain regions were equally affected).

### Reconciling Pser129 pathology with that of previous studies

The OB seeding model we used is associated with impaired odor perception, yet spared motor function (Johnson et al., 2020; Rey et al., 2016). We found, as expected, that injection of α-Syn PFFs into one OB triggered α-Syn accumulation of intraneuronal Pser129 immunoreactive aggregates in both the OB and in inter-connected olfactory structures, particularly in the PCX **(Fig. 4)** which has dense reciprocal connection with the OB. PFF injected mice had significantly greater Pser129 levels compared to mice injected with PBS. Based on prior studies demonstrating Pser129 immunoreactivity in olfactory brain regions of mice following PFF injections (Rey et al., 2018a, 2016), we interpret the Pser129 immunofluorescence we observe in the PCX as being the consequence of seeding of endogenous α-Syn. The uptake pattern we observed is similar to that previously reported (Rey et al., 2016, 2013). Herein and in our earlier studies (Rey et al., 2018a, 2016), injection of PFFs into the OB resulted in elevations in Pser129 relative to the time following seeding. While there was no elevation over time of Pser129 staining in the OB itself, the PCX did show subtle increases in pathology from 1 to 3 months post seeding, albeit not significant. Also similar to our earlier studies (Rey et al., 2016, 2013), we found significantly greater levels of Pser129 in the PCX than the OB.

### α-Syn PFF injection influences odor-evoked beta band activity

Monitoring the spontaneous and odor-evoked LFP activities simultaneously in both the OB and PCX revealed elevations in OB beta-band power in mice with Pser129-positive aggregates. There are several points worthy of discussion in this regard, which we organize by oscillatory band:

### Theta band activity

First, no changes were found in theta-band power, which is a prominent oscillatory power observed in both the OB and PCX. Theta oscillations in the OB are generated by intranasal afferent input, and OB and PCX theta cycles are often coupled to respiration (Kay et al., 2009; Kay and Stopfer, 2006; Komisaruk, 1970). The spared theta power observed herein suggests that OB α-Syn pathology did not overtly alter the basic sensory-motor functions impacting the olfactory system (*i*.*e*., respiration).

### Gamma band activity

Reciprocal dendodendritic activity between OB granule and mitral / tufted cells generates gamma band activity (Shepherd, 1972). Odor-evoked OB gamma oscillations reflect the local network activity within the OB and PCX with gamma in the PCX considered to originate locally (Neville and Haberly, 2003; Rall and Shepherd, 1968). We did not find that PFF injection into the OB affected gamma band activity in either the OB or PCX, during the 3 months post-injection that we followed the mice.

### Beta band activity

Our results indicate that α-Syn aggregates, seeded by PFF injection into the OB, can generate elevations in beta band activity during odor-evoked states **(Fig. 8)**. Beta oscillations reflect large-scale activity between interconnected structures (Kopell et al., 2000; Spitzer and Haegens, 2017) such as that between the OB and PCX (Kay and Beshel, 2010; Neville and Haberly, 2003).

We propose that the changes we uncovered in beta activity are particularly relevant to PD pathophysiology. Striatal and cortical beta band activity is elevated in persons with PD and deep brain stimulation, levodopa, and anti-cholinergic treatment may act by means of suppressing elevated beta power in the cortico-basal ganglia loop (Brown, 2006; Brown et al., 2001; Eisinger et al., 2020; Giannicola et al., 2010; Little and Brown, 2014; McCarthy et al., 2011). Additionally, deep brain stimulation of the subthalamic nucleus was shown in one study to improve cortical function in PD and reduce the excessive beta phase coupling of motor cortex neurons (De Hemptinne et al., 2015). Differences in beta band activity are proposed to be a network determinant of PD pathophysiology (Feingold et al., 2015). Our results indicate an important role for α-Syn related pathologies in these clinical pathophysiologies.

Notably, beta oscillations increase depending upon sensory and cognitive demands (Bauer et al., 2006; Spitzer et al., 2010; Spitzer and Haegens, 2017; van Ede et al., 2010) and in the olfactory system beta oscillations are considered especially involved in odor learning (Gervais et al., 2007; Lowry and Kay, 2007; Martin et al., 2007). In our study, while the mice were awake while being delivered odors, they were not engaged in any task which may influence cognitive demand. Interestingly, in previous work using behavioral tests to assay odor detection and memory, progressive olfactory perceptual deficits, specifically in odor detection and odor retention (memory), were uncovered following injection of PFFs into the OB (Johnson et al., 2020; Rey et al., 2016)., We propose that the aberrant OB odor-evoked beta-band activity we observed is likely to critically influence the PFF-induced changes in odor perception (Johnson et al., 2020; Rey et al., 2016).

We cannot exclude an effect of hormones and biological sex on the influence of PFF injections on beta activity. Work from many labs, including ours, has shown that olfactory system activity can be modulated by sex hormones, including estrogen (Doty and Cameron, 2009; Johnson et al., 2020; Phillips and Vallowe, 1975; Sorwell et al., 2008; Wesson et al., 2006). However, in our study it is unlikely all mice in each group (n>8/group) were at a similar estrous stage during the recordings. Most likely, possible effects of estrous stage would be washed out within groups. Whether or not males injected with PFFs show similar changes in beta activity is an intriguing question, although given the prominence of olfactory dysfunction in male mice following PFF seeding (Johnson et al., 2020), it is likely. Further studies that address the possible influence of biological sex and on olfactory pathophysiology in the context of α-Syn pathology are greatly needed.

### Some brain networks may be more vulnerable to the influence of α-Syn PFF injections than others

The greatest effect of α-Syn PFF injection on neural activity was in the OB. Despite pathogenic propagation of α-Syn from the seeded structure (the OB) into a monosynaptically interconnected structure (the PCX), these structures (*i*.*e*., their networks) were not equally affected. For instance, a striking effect in PFF seeded mice was an elevation in odor-evoked beta power (**Fig. 7 & 8**). This was observed only in the OB – suggesting that the OB is either directly impacted by α-Syn pathology, or, that the PCX, which is known to innervate the OB and influence local inhibition, is impacted and this results in elevated beta power during odor inhalation. We propose the latter model is at play. This is based upon the known nature of beta activity to be originating/communicating between brain regions and also the notably low levels of Pser129 in the OB compared to the PCX. Regardless of mechanism, the results of the odor-evoked analyses highlight that the impact of α-Syn aggregates may be region specific.

In our model, α-Syn aggregates are present within several processing stages of the olfactory system (OB, AON, and PCX). Therefore, we cannot determine with certainty in which neuronal populations (single or multiple) that the α-Syn aggregates influence neural activity, which ultimately give rise to the altered LFPs we detect.

### The influence of α-Syn PFF injection on oscillations is not directly related to levels of pSer129

The altered odor-evoked beta power in PFF injected mice (**Fig. 7 & 8**) was not correlated with levels of pSer129 pathology (**Fig. 9**). The elevations in beta power in the OB during odor were stable regardless of delay post seeding (**Fig. 7 & 8**) which had no impact on pSer129 levels in the OB (**Fig. 4C**).

Cell culture and slice physiology studies demonstrated that pathological α-Syn perturbs normal synaptic function (Volpicelli-Daley et al., 2011; Wu et al., 2019), for instance, by oligomeric α-Syn’s actions upon glutamatergic receptors (Durante et al., 2019). Further, brain-surface electroencephalography from transgenic mice overexpressing human α-synuclein, throughout the brain, uncovered aberrant activity, including epileptiform events (Morris et al., 2015). Our *in vivo* results suggest that elevations in pathological α-Syn, up to a particular level, may be sufficient to entail changes in synaptic coupling or perhaps efficacy which may result in the aberrant LFP activity. While we do not elucidate the precise cellular mechanism whereby α-Syn seeding perturbed neural activity, our correlation results do indicate that function is not strongly correlated with pathogenic α-Syn levels at least at the level of the local networks (*viz*., detectable with LFPs). This outcome points towards the likely influential role of other pathologies related to α-Syn aggregates in shaping neural dynamics in the context of PD. For instance, α-Syn aggregates accumulate in axons where they likely cause degeneration of the axons as well as dendrites (Volpicelli-Daley et al., 2011). Thus, it is possible that the influence of α-Syn aggregates on neural activity would be more directly correlated with changes in activity as the aggregates mature.

### Conclusions

Our results extend important mechanistic work performed in cell culture and in brain slices indicating that α-Syn aggregates can perturb synaptic activity. We used awake mice and studied how progressive changes in α-Syn aggregate pathology affect the olfactory system at the network level. We found a change in neural activity that was region-specific, but did not directly correlate to the local degree of α-Syn aggregate pathology. Future work to assess levels of other pathologies along with simultaneous neural recordings will be informative, as will be work utilizing methods allowing for monitoring the activity of select cell populations to understand the cell types specifically vulnerable to the effects of synucleinopathy.

## Acknowledgements

Research reported in this publication was supported by the National Institute on Deafness and Other Communication Disorders of the National Institutes of Health under Award Numbers R01DC016519 (D.W. and P.B.) and R01DC014443 (D.W.). The content is solely the responsibility of the authors and does not necessarily represent the official views of the National Institutes of Health. We acknowledge the Van Andel Institute and many individuals and corporations that financially support research into neurodegenerative disease at Van Andel Institute. We also thank the Udall Center for Parkinson’s Research at the University of Pennsylvania for generously providing α-Syn PFFs and Karen Kelley at the University of Florida Interdisciplinary Center for Biotechnology Research for expert advice and training in electron microscopy.

## Author contributions

Investigation (A.K., E.R., J.P-B, H.S., M.C.), Formal analyses (A.K., E.R., T.S.), Validation (A.K., T.S., D.W.), Methodology and Resources (J.A.S., K.C.L, and D.W.), Writing-Original draft (A.K.), Review and Editing (A.K., J.A.S., P.B., D.W.), Conceptualization, Supervision, and Funding acquisition (P.B. and D.W.)

## References

Bauer, M., Oostenveld, R., Peeters, M., Fries, P., 2006. Tactile spatial attention enhances gamma-band activity in somatosensory cortex and reduces low-frequency activity in parieto-occipital areas. J. Neurosci. 26, 490–501. https://doi.org/10.1523/JNEUROSCI.5228-04.2006

Braak, H., Ghebremedhin, E., Rüb, U., Bratzke, H., Tredici, K., 2004. Stages in the development of Parkinson’s disease-related pathology. Cell Tissue Res 318, 121–134. https://doi.org/10.1007/s00441-004-0956-9

Braak, H., Rüb, U., Gai, W.P., Del Tredici, K., 2003a. Idiopathic Parkinson’s disease: Possible routes by which vulnerable neuronal types may be subject to neuroinvasion by an unknown pathogen. J. Neural Transm. 110, 517–536. https://doi.org/10.1007/s00702-002-0808-2

Braak, H., Tredici, K., Rüb, U., Vos, R.A.I., Jansen Steur, E.N.H., Braak, E., 2003b. Staging of brain pathology related to sporadic Parkinson’s disease. Neurobiol Aging 24, 197–211. https://doi.org/10.1016/S0197-4580(02)00065-9

Brown, P., 2006. Bad oscillations in Parkinson’s disease, in: Journal of Neural Transmission, Supplement. Springer, Vienna, pp. 27–30. https://doi.org/10.1007/978-3-211-45295-0_6

Brown, P., Oliviero, A., Mazzone, P., Insola, A., Tonali, P., Di Lazzaro, V., 2001. Dopamine Dependency of Oscillations between Subthalamic Nucleus and Pallidum in Parkinson’s Disease. J. Neurosci. 21, 1033–1038. https://doi.org/10.1523/JNEUROSCI.21-03-01033.2001

Brown, P., Williams, D., 2005. Basal ganglia local field potential activity: Character and functional significance in the human. Clin. Neurophysiol. https://doi.org/10.1016/j.clinph.2005.05.009

Burciu, R.G., Vaillancourt, D.E., 2018. Imaging of Motor Cortex Physiology in Parkinson’s Disease. Mov. Disord. 33, 1688–1699. https://doi.org/10.1002/mds.102

Burré, J., 2015. The synaptic function of α-synuclein. J. Parkinsons. Dis. https://doi.org/10.3233/JPD-150642

Burré, J., Sharma, M., Tsetsenis, T., Buchman, V., Etherton, M.R., Südhof, T.C., 2010. α-Synuclein promotes SNARE-complex assembly in vivo and in vitro. Science (80-.). 329, 1663–1667. https://doi.org/10.1126/science.1195227

Buzsáki, G., Anastassiou, C.A., Koch, C., 2012. The origin of extracellular fields and currents — EEG, ECoG, LFP and spikes. Nat Rev Neurosci 13, 407–420. https://doi.org/http://www.nature.com/nrn/journal/v13/n6/suppinfo/nrn3241_S1.html

Chandra, S., Fornai, F., Kwon, H.B., Yazdani, U., Atasoy, D., Liu, X., Hammer, R.E., Battaglia, G., German, D.C., Castillo, P.E., Südhof, T.C., 2004. Double-knockout mice for α- and β-synucleins: Effect on synaptic functions. Proc. Natl. Acad. Sci. U. S. A. 101, 14966–14971. https://doi.org/10.1073/pnas.0406283101

De Hemptinne, C., Swann, N.C., Ostrem, J.L., Ryapolova-Webb, E.S., San Luciano, M., Galifianakis, N.B., Starr, P.A., 2015. Therapeutic deep brain stimulation reduces cortical phase-amplitude coupling in Parkinson’s disease. Nat. Neurosci. 18, 779–786. https://doi.org/10.1038/nn.3997

Doty, R.L., 2017. Olfactory dysfunction in neurodegenerative diseases: is there a common pathological substrate? Lancet Neurol. 16, 478–488. https://doi.org/https://doi.org/10.1016/S1474-4422(17)30123-0

Doty, R.L., 2012. Olfactory dysfunction in Parkinson disease. Nat. Rev. Neurol. 8, 329–339. https://doi.org/10.1038/nrneurol.2012.80

Doty, R.L., Cameron, E.L., 2009. Sex differences and reproductive hormone influences on human odor perception. Physiol. Behav. 97, 213–28. https://doi.org/10.1016/j.physbeh.2009.02.032

Doty, R.L., Ferguson-Segall, M., Lucki, I., Kreider, M., 1988. Effects of intrabulbar injections of 6-hydroxydopamine on ethyl acetate odor detection in castrate and non-castrate male rats. Brain Res 444, 95–103.

Doucette, W., Milder, J., Restrepo, D., 2007. Adrenergic modulation of olfactory bulb circuitry affects odor discrimination. Learn Mem 14, 539–547. https://doi.org/14/8/539 [pii]10.1101/lm.606407

Durante, V., de Iure, A., Loffredo, V., Vaikath, N., De Risi, M., Paciotti, S., Quiroga-Varela, A., Chiasserini, D., Mellone, M., Mazzocchetti, P., Calabrese, V., Campanelli, F., Mechelli, A., Di Filippo, M., Ghiglieri, V., Picconi, B., El-Agnaf, O.M., De Leonibus, E., Gardoni, F., Tozzi, A., Calabresi, P., 2019. Alpha-synuclein targets GluN2A NMDA receptor subunit causing striatal synaptic dysfunction and visuospatial memory alteration. Brain 142, 1365–1385. https://doi.org/10.1093/brain/awz065

Eisinger, R.S., Cagle, J., Opri, E., Alcantara, J., Cernera, S., Foote, K.D., Okun, M.S., Gunduz, A., 2020. Parkinsonian beta dynamics during rest and movement in the dorsal pallidum and subthalamic nucleus. J. Neurosci. 40, 2859–2867. https://doi.org/10.1523/JNEUROSCI.2113-19.2020

Feingold, J., Gibson, D.J., DePasquale, B., Graybiel, A.M., 2015. Bursts of beta oscillation differentiate postperformance activity in the striatum and motor cortex of monkeys performing movement tasks. Proc. Natl. Acad. Sci. 112, 13687 LP – 13692. https://doi.org/10.1073/pnas.1517629112

Fujiwara, H., Hasegawa, M., Dohmae, N., Kawashima, A., Masliah, E., Goldberg, M.S., Shen, J., Takio, K., Iwatsubo, T., 2002. α-Synuclein is phosphorylated in synucleinopathy lesions. Nat. Cell Biol. 4, 160–164. https://doi.org/10.1038/ncb748

Gadziola, M.A., Tylicki, K.A., Christian, D.L., Wesson, D.W., 2015. The Olfactory Tubercle Encodes Odor Valence in Behaving Mice. J. Neurosci. 35, 4515–4527. https://doi.org/10.1523/JNEUROSCI.4750-14.2015

Gervais, R., Buonviso, N., Martin, C., Ravel, N., 2007. What do electrophysiological studies tell us about processing at the olfactory bulb level? J. Physiol. Paris 101, 40–45. https://doi.org/10.1016/j.jphysparis.2007.10.006

Giannicola, G., Marceglia, S., Rossi, L., Mrakic-Sposta, S., Rampini, P., Tamma, F., Cogiamanian, F., Barbieri, S., Priori, A., 2010. The effects of levodopa and ongoing deep brain stimulation on subthalamic beta oscillations in Parkinson’s disease. Exp. Neurol. 226, 120–127. https://doi.org/10.1016/j.expneurol.2010.08.011

Goedert, M., 2001. Alpha-synuclein and neurodegenerative diseases. Nat Rev Neurosci 2. https://doi.org/10.1038/35081564

Gottfried, J.A., 2010. Central mechanisms of odour object perception. Nat Rev Neurosci 11, 628–641. https://doi.org/http://www.nature.com/nrn/journal/v11/n9/suppinfo/nrn2883_S1.html

Haegens, S., Zion Golumbic, E., 2018. Rhythmic facilitation of sensory processing: A critical review. Neurosci. Biobehav. Rev. 86, 150–165. https://doi.org/https://doi.org/10.1016/j.neubiorev.2017.12.002

Hammond, C., Bergman, H., Brown, P., 2007. Pathological synchronization in Parkinson’s disease: networks, models and treatments. Trends Neurosci. https://doi.org/10.1016/j.tins.2007.05.004

Johnson, M.E., Bergkvist, L., Mercado, G., Stetzik, L., Meyerdirk, L., Wolfrum, E., Madaj, Z., Brundin, P., Wesson, D., 2020. Deficits in olfactory sensitivity in a mouse model of Parkinson’s disease revealed by plethysmography of odor-evoked sniffing. Sci. Rep. 10, 1–27. https://doi.org/10.1101/2020.02.27.968545

Kay, L.M., Beshel, J., 2010. A beta oscillation network in the rat olfactory system during a 2-alternative choice odor discrimination task. J. Neurophysiol. 104, 829–839.

Kay, L.M., Beshel, J., Brea, J., Martin, C., Rojas-Líbano, D., Kopell, N., 2009. Olfactory oscillations: the what, how and what for. Trends Neurosci. 32, 207–214.

Kay, L.M., Stopfer, M., 2006. Information processing in the olfactory systems of insects and vertebrates. Semin. Cell Dev. Biol. 17, 433–442. https://doi.org/http://dx.doi.org/10.1016/j.semcdb.2006.04.012

Komisaruk, B.R., 1970. Synchrony between limbic system theta activity and rhythmical behavior in rats. J Comp Physiol Psychol 70, 482–492.

Kopell, N., Ermentrout, G.B., Whittington, M.A., Traub, R.D., 2000. Gamma rhythms and beta rhythms have different synchronization properties. Proc. Natl. Acad. Sci. U. S. A. 97, 1867–1872.

Kühn, A.A., Kupsch, A., Schneider, G.H., Brown, P., 2006. Reduction in subthalamic 8-35 Hz oscillatory activity correlates with clinical improvement in Parkinson’s disease. Eur. J. Neurosci. 23, 1956–1960. https://doi.org/10.1111/j.1460-9568.2006.04717.x

Lee, E.Y., Eslinger, P.J., Du, G., Kong, L., Lewis, M.M., Huang, X., 2014. Olfactory-related cortical atrophy is associated with olfactory dysfunction in Parkinson’s disease. Mov. Disord. 29, 1205–1208. https://doi.org/10.1002/mds.25829

Little, S., Brown, P., 2014. The functional role of beta oscillations in Parkinson’s disease. Park. Relat. Disord. 20, S44–S48. https://doi.org/10.1016/S1353-8020(13)70013-0

Lowry, C.A., Kay, L.M., 2007. Chemical factors determine olfactory system beta oscillations in waking rats. J Neurophysiol 98, 394–404. https://doi.org/00124.2007 [pii]10.1152/jn.00124.2007

Luk, K., Kehm, V., Zhang, B., O’Brien, P., Trojanowski, J., Lee, V., 2012. Intracerebral inoculation of pathological α-synuclein initiates a rapidly progressive neurodegenerative α-synucleinopathy in mice. J Exp Med 209. https://doi.org/10.1084/jem.20112457

Luk, K.C., Lee, V.M.-Y., 2014. Modeling Lewy pathology propagation in Parkinson’s disease. Parkinsonism Relat. Disord. 20, S85–S87. https://doi.org/10.1016/S1353-8020(13)70022-1

Luk, K.C., Song, C., O’Brien, P., Stieber, A., Branch, J.R., Brunden, K.R., Trojanowski, J.Q., Lee, V.M.Y., 2009. Exogenous α-synuclein fibrils seed the formation of Lewy body-like intracellular inclusions in cultured cells. Proc. Natl. Acad. Sci. U. S. A. 106, 20051–20056. https://doi.org/10.1073/pnas.0908005106

Mahlknecht, P., Seppi, K., Poewe, W., 2015. The concept of prodromal Parkinson’s disease. J. Parkinsons. Dis. https://doi.org/10.3233/JPD-150685

Martin, C., Beshel, J., Kay, L.M., 2007. An olfacto-hippocampal network is dynamically involved in odor-discrimination learning. J Neurophysiol 98, 2196–2205.

Mason, D.M., Nouraei, N., Pant, D.B., Miner, K.M., Hutchison, D.F., Luk, K.C., Stolz, J.F., Leak, R.K., 2016. Transmission of α-synucleinopathy from olfactory structures deep into the temporal lobe. Mol. Neurodegener. 11, 49. https://doi.org/10.1186/s13024-016-0113-4

McCarthy, M.M., Moore-Kochlacs, C., Gu, X., Boyden, E.S., Han, X., Kopell, N., 2011. Striatal origin of the pathologic beta oscillations in Parkinson’s disease. Proc. Natl. Acad. Sci. U. S. A. 108, 11620–11625. https://doi.org/10.1073/pnas.1107748108

Mezias, C., Rey, N., Brundin, P., Raj, A., 2020. Neural connectivity predicts spreading of alpha-synuclein pathology in fibril-injected mouse models: Involvement of retrograde and anterograde axonal propagation. Neurobiol. Dis. 134, 104623. https://doi.org/10.1016/j.nbd.2019.104623

Mitzdorf, U., 1985. Current source-density method and application in cat cerebral cortex: investigation of evoked potentials and EEG phenomena. Physiol. Rev. 65, 37–100.

Moran, A., Bergman, H., Israel, Z., Bar-Gad, I., 2008. Subthalamic nucleus functional organization revealed by parkinsonian neuronal oscillations and synchrony. Brain 131, 3395–409. https://doi.org/10.1093/brain/awn270

Morris, M., Sanchez, P.E., Verret, L., Beagle, A.J., Guo, W., Dubal, D., Ranasinghe, K.G., Koyama, A., Ho, K., Yu, G.Q., Vossel, K.A., Mucke, L., 2015. Network dysfunction in α-synuclein transgenic mice and human Lewy body dementia. Ann. Clin. Transl. Neurol. 2, 1012–1028. https://doi.org/10.1002/acn3.257

Neville, K.R., Haberly, L.B., 2003. Beta and gamma oscillations in the olfactory system of the urethane-anesthetized rat. J Neurophysiol 90, 3921–3930.

Nusser, Z., Kay, L.M., Laurent, G., Homanics, G.E., Mody, I., 2001. Disruption of GABA(A) receptors on GABAergic interneurons leads to increased oscillatory power in the olfactory bulb network. J Neurophysiol 86, 2823–2833.

Paxinos, G., Franklin, K., 2000. The Mouse Brain in Stereotaxic Coordinates, 2nd ed. Academic Press, San Diego.

Peelaerts, W., Bousset, L., Baekelandt, V., Melki, R., 2018. α-Synuclein strains and seeding in Parkinson’s disease, incidental Lewy body disease, dementia with Lewy bodies and multiple system atrophy: similarities and differences. Cell Tissue Res. https://doi.org/10.1007/s00441-018-2839-5

Phillips, P.D., Vallowe, H.H., 1975. Cyclic fluctuations in odor detection by female rats and the temporal influences of exogenous steroids on ovariectomized rats. Proc. Pennsylvania Acad. Sci. https://doi.org/10.2307/44110934

Polinski, N.K.., Volpicelli-Daley, L.A.., Sortwell, C.E.., Luk, K.C.., Cremades, N., Gottler, L.M.., Froula, J., Duffy, M.F.., Lee, V.M.., Martinez, T.N.., Dave, K.D.., 2018. Best Practices for Generating and Using Alpha-Synuclein Pre-Formed Fibrils to Model Parkinson’s Disease in Rodents. J. Parkinsons. Dis.

Rall, W., Shepherd, G.M., 1968. Theoretical reconstruction of field potentials and dendrodendritic synaptic interactions in olfactory bulb. J Neurophysiol 31, 884–915.

Rey, N.L., George, S., Steiner, J.A., Madaj, Z., Luk, K.C., Trojanowski, J.Q., Lee, V.M.-Y., Brundin, P., 2018a. Spread of aggregates after olfactory bulb injection of α-synuclein fibrils is associated with early neuronal loss and is reduced long term. Acta Neuropathol. 135, 65–83. https://doi.org/10.1007/s00401-017-1792-9

Rey, N.L., Petit, G.H., Bousset, L., Melki, R., Brundin, P., 2013. Transfer of human α-synuclein from the olfactory bulb to interconnected brain regions in mice. Acta Neuropathol. 126, 555–573. https://doi.org/10.1007/s00401-013-1160-3

Rey, N.L., Steiner, J.A., Maroof, N., Luk, K.C., Madaj, Z., Trojanowski, J.Q., Lee, V.M.-Y., Brundin, P., 2016. Widespread transneuronal propagation of α-synucleinopathy triggered in olfactory bulb mimics prodromal Parkinson’s disease. J. Exp. Med. 213, 1759–78. https://doi.org/10.1084/jem.20160368

Rey, N.L., Wesson, D.W., Brundin, P., 2018b. The olfactory bulb as the entry site for prion-like propagation in neurodegenerative diseases. Neurobiol. Dis. 109, 226–248. https://doi.org/http://dx.doi.org/10.1016/j.nbd.2016.12.013

Ross, G.W., Petrovitch, H., Abbott, R.D., Tanner, C.M., Popper, J., Masaki, K., Launer, L., White, L.R., 2008. Association of olfactory dysfunction with risk for future Parkinson’s disease. Ann. Neurol. 63, 167–173. https://doi.org/10.1002/ana.21291

Schindelin, J., Arganda-Carreras, I., Frise, E., Kaynig, V., Longair, M., Pietzsch, T., Preibisch, S., Rueden, C., Saalfeld, S., Schmid, B., Tinevez, J.Y., White, D.J., Hartenstein, V., Eliceiri, K., Tomancak, P., Cardona, A., 2012. Fiji: An open-source platform for biological-image analysis. Nat. Methods. https://doi.org/10.1038/nmeth.2019

Schoppa, N.E., Urban, N.N., 2003. Dendritic processing within olfactory bulb circuits. Trends Neurosci 26, 501–506.

Scott, J.W., McBride, R.L., Schneider, S.P., 1980. The organization of projections from the olfactory bulb to the piriform cortex and olfactory tubercle in the rat. J Comp Neurol 194, 519–534.

Shepherd, G.M., 1972. Synaptic organization of the mammalian olfactory bulb. Physiol. Rev. https://doi.org/10.1152/physrev.1972.52.4.864

Sorwell, K.G., Wesson, D.W., Baum, M.J., 2008. Sexually dimorphic enhancement by estradiol of male urinary odor detection thresholds in mice. Behav. Neurosci. 122, 788–793. https://doi.org/2008-09788-007 [pii] 10.1037/0735-7044.122.4.788

Spillantini, M.G., Schmidt, M.L., Lee, V.M., Trojanowski, J.Q., Jakes, R., Goedert, M., 1997. Alpha-synuclein in Lewy bodies. Nature 388. https://doi.org/10.1038/42166

Spitzer, B., Haegens, S., 2017. Beyond the Status Quo: A Role for Beta Oscillations in Endogenous Content (Re)Activation. eneuro 4, ENEURO.0170-17.2017. https://doi.org/10.1523/ENEURO.0170-17.2017

Spitzer, B., Wacker, E., Blankenburg, F., 2010. Oscillatory correlates of vibrotactile frequency processing in human working memory. J. Neurosci. 30, 4496–4502. https://doi.org/10.1523/JNEUROSCI.6041-09.2010

van Ede, F., Jensen, O., Maris, E., 2010. Tactile expectation modulates pre-stimulus A-band oscillations in human sensorimotor cortex. Neuroimage 51, 867–876. https://doi.org/10.1016/j.neuroimage.2010.02.053

Volpicelli-Daley, L.A., Luk, K.C., Lee, V.M.-Y., 2014. Addition of exogenous α-synuclein preformed fibrils to primary neuronal cultures to seed recruitment of endogenous α-synuclein to Lewy body and Lewy neurite–like aggregates. Nat. Protoc. 9, 2135.

Volpicelli-Daley, L.A., Luk, K.C., Patel, T.P., Tanik, S.A., Riddle, D.M., Stieber, A., Meaney, D.F., Trojanowski, J.Q., Lee, V.M.-Y., 2011. Exogenous α-synuclein fibrils induce Lewy body pathology leading to synaptic dysfunction and neuron death. Neuron 72, 57–71. https://doi.org/10.1016/j.neuron.2011.08.033

Wachowiak, M., Shipley, M.T., 2006. Coding and synaptic processing of sensory information in the glomerular layer of the olfactory bulb. Semin Cell Dev Biol 17, 411–423.

Wattendorf, E., Welge-Lussen, A., Fiedler, K., Bilecen, D., Wolfensberger, M., Fuhr, P., Hummel, T., Westermann, B., 2009. Olfactory Impairment Predicts Brain Atrophy in Parkinson’s Disease. J. Neurosci. 29, 15410–15413. https://doi.org/10.1523/jneurosci.1909-09.2009

Wen, M.C., Xu, Z., Lu, Z., Chan, L.L., Tan, E.K., Tan, L.C.S., 2017. Microstructural network alterations of olfactory dysfunction in newly diagnosed Parkinson’s disease. Sci. Rep. 7, 12559. https://doi.org/10.1038/s41598-017-12947-7

Wesson, D.W., Keller, M., Douhard, Q., Baum, M.J., Bakker, J., 2006. Enhanced urinary odor discrimination in female aromatase knockout (ArKO) mice. Horm. Behav. 49, 580–586.

Wilson, D.A., 2001. Scopolamine enhances generalization between odor representations in rat olfactory cortex. Learn Mem 8, 279–285.

Wilson, D.A., Sullivan, R.M., 2011. Cortical processing of odor objects. Neuron 72, 506–519.

Wu, Q., Takano, H., Riddle, D.M., Trojanowski, J.Q., Coulter, D.A., Lee, V.M.-Y., 2019. Alpha-synuclein (αSyn) preformed fibrils induce endogenous αSyn aggregation, compromise synaptic activity and enhance synapse loss in cultured excitatory hippocampal neurons. J. Neurosci. 0060–19. https://doi.org/10.1523/JNEUROSCI.0060-19.2019

Wu, X., Yu, C., Fan, F., Zhang, K., Zhu, C., Wu, T., Li, K., Chan, P., 2011. Correlation between Progressive Changes in Piriform Cortex and Olfactory Performance in Early Parkinson’s Disease. Eur. Neurol. 66, 98–105. https://doi.org/10.1159/000329371

